# Valosin-containing Protein is Cargo in Amyloid Precursor Protein Extracellular Vesicles

**DOI:** 10.1101/2025.01.20.633888

**Authors:** Yue Lu, Mohammad Abdullah, Liam R. Healy, Marc D. Tambini

## Abstract

The Amyloid Precursor Protein (APP), a genetic cause of Alzheimer’s disease (AD), is a type-I transmembrane protein that is metabolized by proteolysis in the endolysomal system. APP and its metabolites are secreted by cells in extracellular vesicles (EVs). To study the function of APP-containing EVs, we isolated App-EVs from rat primary neuronal conditioned media and proteomic analysis identified the Valosin-containing protein (Vcp) as molecular cargo. Pharmacological modulation of Vcp activity was found to alter App processing and global EV secretion in rat primary neurons. AD-associated knock-in *App* mutations were found to alter the abundance of App-EVs and the trafficking of App metabolites within App-EVs, in a manner related to the epitopes generated by the nonamyloidogenic processing of App. The presence of Vcp suggests a role for App-EVs in the clearance of protein aggregates.

## INTRODUCTION

Alzheimer’s disease (AD) is the most common form of dementia in the elderly and a major cause of morbidity and mortality worldwide^1^. It is characterized by progressive neuronal loss and the histopathological appearance of two canonical lesions: extracellular senile plaques, composed of aggregated amyloid beta (Aβ), and intracellular neurofibrillary tangles, composed of hyperphosphorylated tau^2^. Mutations in the *Amyloid Precursor Protein* (*APP*-human, *App* - rodent), a type-I transmembrane protein that undergoes sequential proteolysis to produce Aβ, cause familial forms of AD^3^. Recently, three anti-Aβ antibodies have been shown to effectively reduce senile plaque deposits^4–6^, but, despite this reduction, AD patients will still invariably develop dementia. An understanding of the pathogenic effects of APP and Aβ beyond plaque formation is therefore needed.

APP undergoes extensive intracellular processing which results in multiple APP metabolites^3^. The majority of APP processing occurs via the nonamyloidogenic pathway, in which APP is initially cleaved in the juxtamembranous region by α-secretase to produce a large soluble ectodomain (sAPPα), and the membrane bound C-terminal fragment (α-CTF). APP α-CTF undergoes further processing by γ-secretase to release the non-aggregating p3 peptide and an APP intracellular domain (AICD). Amyloidogenic processing of APP begins with β-secretase cleavage to produce a soluble APP ectodomain (sAPPβ) and a membrane bound C-terminal fragment (β-CTF). APP β-CTF is further cleaved by γ-secretase to release Aβ and AICD. The length of Aβ can vary and determines its propensity to aggregate, with shorter forms, such as Aβ40, less likely to aggregate and longer forms, such as Aβ42 and longer, more likely to aggregate. Given the numerous APP metabolites that neurons produce, the typical changes (〈 total Aβ, 〈Aβ42:40 ratio) that track with plaque formation do not adequately capture the full spectrum of APP metabolism. Moreover, familial *APP* mutations that increase amyloidogenic processing of APP also result in the dysregulation of these APP metabolites, many of which exert functions within the cell independent of plaque pathology.

Numerous lines of evidence support a role for APP and its metabolites in the endolysosomal system, whose dysfunction is an early pathological change in AD^7–9^. While APP is initially trafficked along the biosynthetic-secretory pathway, its processing occurs at the plasma membrane^10^ (α- and γ-cleavage) and within the endolysosomal system^11^ (β- and γ-cleavage). Multiple APP metabolites have been found to accumulate in the endolysosomes, and, in particular, within multivesicular bodies^12–14^ (MVBs). MVBs are formed when the outer, or limiting, membrane of endosomes invaginates to form intralumenal vesicles (ILVs). Upon fusion of the MVB to the plasma membrane, ILVs are released into the extracellular space where they can be taken up or exert effects on distant cells. These secreted ILVs are termed exosomes, a subset of extracellular vesicles (EVs). EVs have been evaluated for their role in the production and spread of Aβ^15^, though it has also been observed that other APP metabolites, such as APP-CTFs, selectively accumulate in EVs as well^16–19^. The function of these APP-containing EVs (APP-EVs - human, App-EVs - rodent) is unknown. Several technical limitations contribute to this lack of understanding: 1. All cells secrete EVs, and it can be difficult to determine the origin of EVs derived from a source containing multiple cell types, such as in the brain^20^. 2. Within one cell, multiple different pathways result in ILV/EV formation. Invagination of the limiting membrane of endosomes occurs at different places in the cell and in response to different signals, resulting in the secretion of a heterogenous mixture of EVs with varying cargoes^21^. 3. EV isolation methods rely on the bulk purification of total EVs from this heterogeneous population, or the selective enrichment of EVs based on candidate markers. However, as with cargoes, EV markers are not present in every EV and therefore define only a subset of the total population^22,23^.

Here, we report the immunocapture and analysis of purified App-EVs derived from rat primary neuronal conditioned media. We find that App-EVs contain as molecular cargo the valosin-containing protein (Vcp), a ubiquitin-dependent segregase/molecular unfoldase and the genetic cause of autosomal dominant forms of AD related dementias^24^. We uncover a new ability of Vcp to regulate App metabolism and global EV secretion in primary neurons. Using a genetically faithful rat knock-in model of an AD-associated *App* mutation, we link App-EV biogenesis to the nonamyloidogenic App processing pathway. Together, these results point to a new function of App and its processing that may relate to the clearance of aggregated proteins via Vcp-containing App-EVs.

## RESULTS

### App-EVs contain Vcp

To understand the function of App-EVs it is necessary to isolate App-EVs from other vesicles. This presents several considerations, such as cell-type source, organism source, and purification method. App is expressed in multiple cell types, but given that App is predominantly expressed in neurons^25^, and that neuronal cell death underlies the progressive cognitive impairment seen in AD, the choice of neurons is most relevant. EV isolation from primary neuronal conditioned media allows for the study of exclusively neuronal EVs and removes the possibility of contamination with intracellular vesicles, which have the same biophysical properties, such as size and density, and protein markers as EVs. For the study of AD, it would be ideal to use a human source of neurons, but given the scale of induced pluripotent stem cell (iPSC)-derived neurons required for App-EV isolation and the nonphysiological features of neuronal cancer cell lines, the ability to study neuronal App-EVs from human sources is limited. Transgenic animals have been engineered to express human *APP,* with and without AD-associated mutations, though many rodent models rely on overexpression of the transgene or the use of multiple AD-mutations, both of which may alter the physiological function of APP^26^. Recently, *App* knock-in rats (*App^h^* and *App^S^*) have been developed which offer advantages over other transgenic animals^27–29^. *App^h^* rats express a humanized form of rodent *App* under the control of the endogenous rodent *App* promoter, therefore, each *App* rat in this study produces human Aβ, human p3, and human App-CTFs. To examine the effect of AD-related amyloidogenic App processing on App-EV function, *App^S^* rats were engineered to additionally express the Swedish mutation, which drives *App* metabolism toward amyloidogenic processing in a manner similar to familial AD patients with the Swedish *APP* mutation. For these technical considerations, conditioned media from primary neurons from *App^h^* and *App^S^*rats were used in this study.

Immunocapture of EVs using antibodies against common EV markers such as Alix, CD9, CD63, or CD81 has been found to enrich EVs from a heterogenous mixture of vesicles^23^, and, as *App* is a type-I transmembrane protein with its N-terminus exposed to the extracellular space in EVs, we predicted that App-EVs could be isolated by similar methods using anti-App antibodies. Neuronal conditioned media was filtered to remove large debris and concentrated 80X for use as input. Concentration of EV-containing media was chosen over the standard use of ultracentrifugation to pellet EVs because ultracentrifugation has been shown to cause aggregation of vesicles^30^ which may result in nonspecific co-immunoprecipitation of non-App-EVs. The anti-App 4G8 antibody was chosen, as it recognizes epitopes at the juxtamembranous extracellular-facing region of App and does not recognize, and therefore does not compete against, the abundant App-metabolite sAppα present in conditioned media (**Fig. 1A**). The detection of ∼110 kDa App with a C-terminal anti-App antibody in anti-App 4G8 eluate indicates the presence of the full length protein, including its transmembrane domain (**Fig. 1B**). As no detergents were used in the immunocapture, the presence of the full length App protein suggests a membranous source, which we term App-EVs. Mass spectrometry sequencing of immunoisolated App-EVs from *App^h^* and *App^S^* neuronal conditioned media revealed the presence of Vcp in both samples, with no Vcp peptides detected in IgG controls (**Fig. 1C**). Vcp is an abundant multifunctional protein that binds and unfolds multiple protein substrates, including polyubiquitinated aggregates, for degradation^24^. In addition to Vcp, known Vcp interactors, including polyubiquitin^31^, histone subunits^32,33^, and ribosome subunits^32,34,35^, were detected (**Fig. 1C**). Western analysis of App-EVs confirmed the presence of Vcp in *App^h^* and *App^S^* samples (**Fig. 1D**). Direct binding of App and Vcp was investigated by co-immunoprecipitation of App and Vcp in Triton-X solubilized brain lysate (**Fig. S1**). The absence of co-immunoprecipitation of App by Vcp, and vice-versa, suggests a lack of direct binding and excludes the possibility that Vcp is binding aggregated App. App-EVs were further characterized by western analysis using antibodies against the common EV markers flotillin-1, Alix, and CD9 (**Fig. 1E**). These markers were not detected in App-EV samples, in agreement with their absence from the mass spectrometric results. Given that these common EV markers were not present in App-EVs, further confirmation of the App-EV’s vesicular identity was accomplished using transmission electron microscopic analysis (**Fig. 1F**). Numerous spherical particles less than 40 nm in diameter were detected and are consistent with electron micrographs of small EVs^36^. The combined immunoprecipitation, mass spectrometric, and electron micrographic results support the conclusion that App-EVs have been isolated.

**Figure 1.**
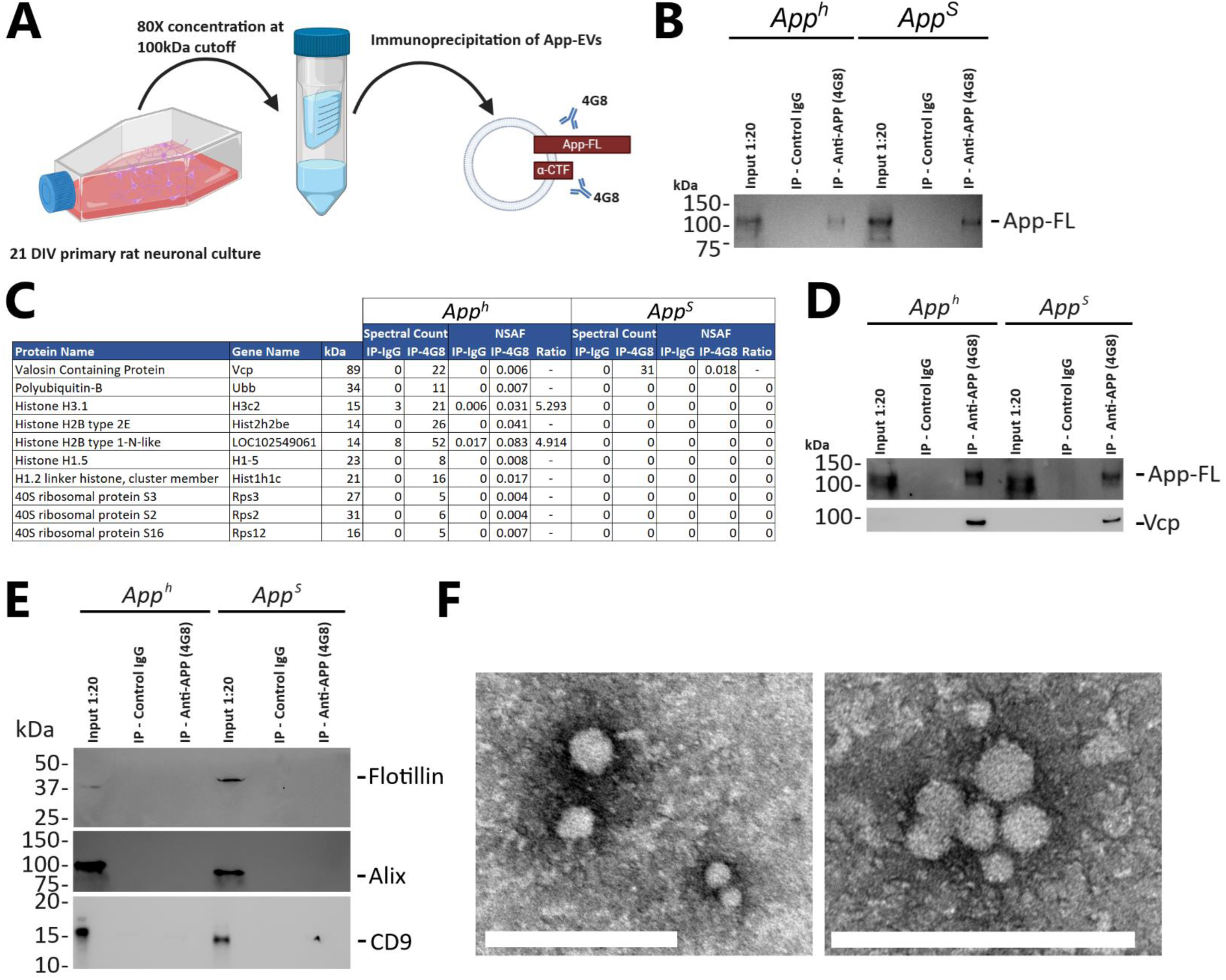
Isolation and characterization of App-EVs. **A**. Schematic of App-EV isolation from filtered and concentrated *App^h^* and *App^S^* rat primary neuronal conditioned media, using anti-App 4G8 antibody directed against the extracellular-facing juxtamembranous region of App full length and App-CTFs. **B.** Immunocapture of App-EVs with 4G8 or control IgG, followed by western analysis with anti-App Y188 antibody directed against the C-terminus of App. Input samples were diluted 20X. **C.** Mass spectrometry analysis of App-EVs immunoisolated from primary neuronal conditioned media vs control IgG. Spectral Counts and Normalized Spectral Abundance Factors are shown. **D.** Co-immunoprecipitation of Vcp in immunocaptured App-EVs. App-EVs were immunoisolated by 4G8 or control IgG, followed by western analysis with anti-Vcp antibody. Input samples were diluted 20X. n=3. **E.** App-EVs were immunoisolated by 4G8 or control IgG, followed by western analysis with flotillin-1, Alix, and CD9. Input samples were diluted 20X. **F.** TEM images of immunoisolated App-EVs at 100k X (left) and 150k X (right). Scale bars indicate 200 nm.

### Vcp inhibition causes global EV release

Given the finding that App-EVs contain Vcp cargo, we next investigated the functional effect of Vcp on App levels and metabolism. While *Vcp* knockout is embryonically lethal^37^, pharmacological inhibition of Vcp can be accomplished by NMS-873, a potent allosteric inhibitor which binds the region between the Vcp D1 and D2 ATPase domains^38^. 8h treatment of primary *App^h^* neurons with 2.5 μM NMS-873 resulted in significantly less App α-CTF in cell lysate, with no concomitant changes to full length App levels (**Fig. 2A**). This result was mirrored by the increase in App α-CTF caused by the dual activation of D1 and D2 domains by Smer28^39^ and VA1^40^, respectively (**Fig. S2**). The effect of Vcp inhibition on App α-CTF independent of full length App suggests that lower App α-CTF levels are not the result of a transcriptional response. Decreased production (by reduced α-secretase activity) or increased clearance (by increased macroautophagy or γ-secretase activity) could explain lower App α-CTF levels. sAppα and App α-CTF are produced in equimolar amounts when App is cleaved by α-secretase; therefore, sAppα levels in conditioned media indicate α-secretase activity. Paradoxically, increased sAppα levels were observed in NMS-873-treated samples (**Fig. 2B**), which rules out a decrease in production. We next focused on the effect of NMS-873 on major App-CTF degradative pathways, including macroautophagy and γ-secretase proteolysis. Vcp is required for autophagy and binds Beclin 1 and the PI3K complex^41,42^. In agreement with this function, we find that, rather than increasing autophagy, inhibition of Vcp by NMS-873 lowers autophagy, as indicated by a lower LC3II/I ratio in chloroquine-treated samples (**Fig. 2C**). γ-Secretase activity results in the production of Aβ, which can be measured in neuronal conditioned media. No significant difference in the most abundant form of Aβ, Aβ40, was observed in NMS-873-treated samples, while a statistically significant decrease in the second most abundant form, Aβ42, was detected (**Fig. 2D**). Together, these data indicate that neither decreased production nor increased clearance is responsible for the NMS-873-mediated decrease in App α-CTF.

**Figure 2.**
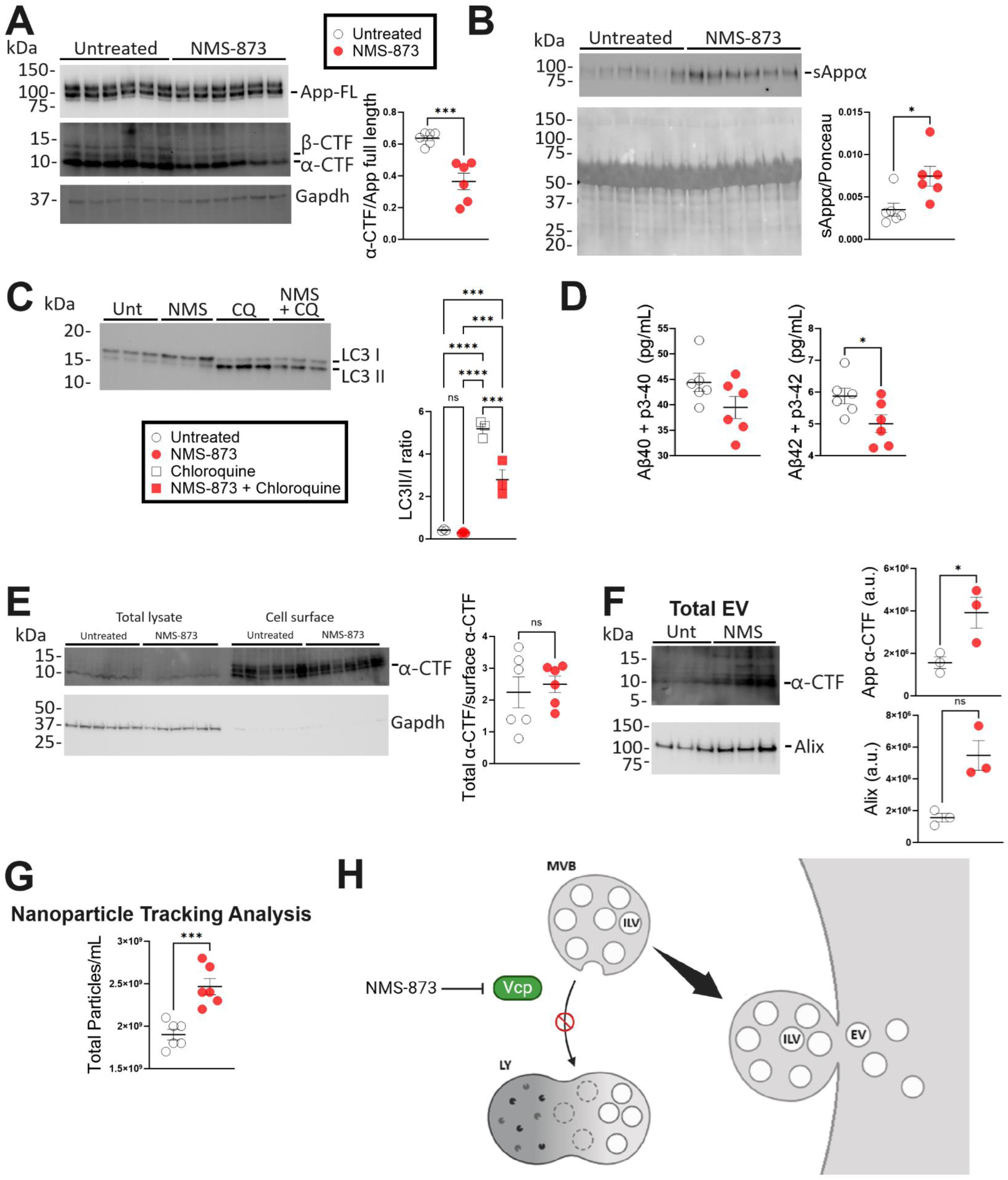
Effect of Vcp inhibition on App processing and EV release. **A**. Western analysis of *App^h^* primary neurons treated with 2.5 μM NMS-873 for 8h. App full length, App β- and α-CTFs, and Gapdh are indicated. Levels of App α-CTF relative to App full length are represented as mean ± S.E.M. and were analyzed by Student’s *t*-test. \*\*\**p* < .001, n=6. **B.** Western analysis of conditioned media from NMS-873-treated *App^h^* primary neurons. sAppα was detected with anti-App 6E10 directed against the Aβ 3-8 region. Red Ponceau is shown below western blot. Levels of sAppα relative to Ponceau stain are represented as mean ± S.E.M. and were analyzed by Student’s *t*-test. \**p* < .05, n=6. **C.** Western analysis of *App^h^* primary neurons treated with 2.5 μM NMS-873 and/or 50 μM chloroquine for 8h. LC3 I and LC3 II were detected with an antibody against LC3B. LC3 II/I ratios are represented as mean ± S.E.M. and were analyzed by one-way ANOVA with Tukey’s multiple comparison test when ANOVA showed significant differences. \*\*\**p* < .001, \*\*\*\**p* < .0001, n=3. **D.** MSD electrochemiluminescent assay of conditioned media from NMS-873-treated *App^h^* primary neurons. Aβ40 and p3-40 were detected with a capture antibody against the C-terminus of Aβ40 and a 4G8 detection antibody. In the same well, Aβ42 and p3-42 were detected with a capture antibody against the C-terminus of Aβ42 and a 4G8 detection antibody. Aβ and p3 levels are represented as mean ± S.E.M and were analyzed by Student’s *t*- test. \**p* < .05, n=6. **E.** Western analysis of total (left) and cell-surface (right) protein levels of *App^h^* primary neurons treated with 2.5 μM NMS-873 for 8h. App α-CTF and Gapdh are indicated. Total App α-CTF levels normalized to surface App α-CTF are represented as mean ± S.E.M and were analyzed by Student’s *t*-test, n=6. **F.** Western analysis of total EVs from conditioned media of NMS-873-treated *App^h^* primary neurons. Alix and App α-CTF are indicated. App α-CTF and Alix levels are represented as mean ± S.E.M. and were analyzed by Student’s *t*-test. \**p* < .05, n=3. **G.** Nanoparticle tracking analysis of conditioned media from *App^h^* primary neurons treated with NMS-873. Total particle levels are represented as mean ± S.E.M. and were analyzed by Student’s *t*-test. \*\*\**p* < .001, n=6. **H.** Summary schematic of NMS-873-induced EV release.

The significant increase in sAppα (**Fig. 2B**) suggests an increased amount of App is trafficked to the cell surface, the predominant subcellular localization of α-secretase activity. Therefore, we used a cell-surface labelling assay to determine cell surface levels of App α-CTF in NMS-873-treated neurons. Despite a decrease in total App α-CTF levels, no such difference was observed at the cell surface (**Fig. 2E**), implicating a change in App trafficking. One possible change in App trafficking that would result in more App available for α- secretase processing at the cell surface is the fusion of App-containing MVBs to the cell surface. The ILVs within, which are selectively enriched in App-CTFs^16–19^, would be secreted and detected in total EV preparations. Increased MVB fusion to the cell surface would likewise result in decreased cellular levels of App-CTFs (**Fig. 2A**). Western analysis of Alix and App α-CTF showed significantly increased levels in conditioned media from NMS-873-treated neurons, consistent with increased EV secretion (**Fig. 2F**). Increased global EV secretion was confirmed with nanoparticle tracking analysis (**Fig. 2G**). We hypothesize that the reduction of autophagy by Vcp inhibitors results in the secretion of EVs, a phenomenon which has been seen with other modulators of autophagy^43,44^ (**Fig. 2H**).

### *App^S^* mutation reduces App-EV levels

App is extensively processed by sequential proteolysis along the nonamyloidogenic and amyloidogenic pathways. These pathways result in different App membrane-bound CTFs which have been reported to be enriched in App-EVs^16–19^. The use of genetic AD-associated mutants would allow us to determine the effect of alterations in App processing on App-EV composition and function. One well-characterized *App* mutation, the Swedish mutation, is located at the two amino acids N-terminal to the β-cleavage site^45^. Swedish-App preferentially binds β-secretase and commits App to the amyloidogenic pathway, generating App β-CTF and, in turn, Aβ. Wild type App is processed primarily along the nonamyloidogenic pathway, which results in App α-CTF, and, in turn, p3. The Swedish mutation-induced shift toward amyloidogenic App processing is recapitulated in *App^S^* rat primary neurons, which display significantly higher App β-CTF levels than *App^h^* controls, where β-CTF is often below the limits of detection (**Fig. 3A**). To determine if the cellular changes in App metabolite abundance caused by the Swedish mutation are reflected in EVs, we isolated total EVs from *App^h^* and *App^S^* primary neuronal conditioned media and analyzed App content by western blot. Significant decreases in both full length App and App α-CTF were observed in *App^S^*samples (**Fig. 3B**). App β-CTF, which composed roughly half of total App-CTFs in *App^S^* neuronal lysates, was strikingly undetectable in EVs, and therefore omitted from quantifications where relevant. To determine if this decrease in full length App and App-CTFs was the result of a global reduction in EVs, conditioned media from *App^h^* and *App^S^*primary neuronal cultures was analyzed by nanoparticle tracking, which showed no differences (**Fig. 3C**). The effect of the Swedish *App* mutation appears to be confined to App-EVs.

**Figure 3.**
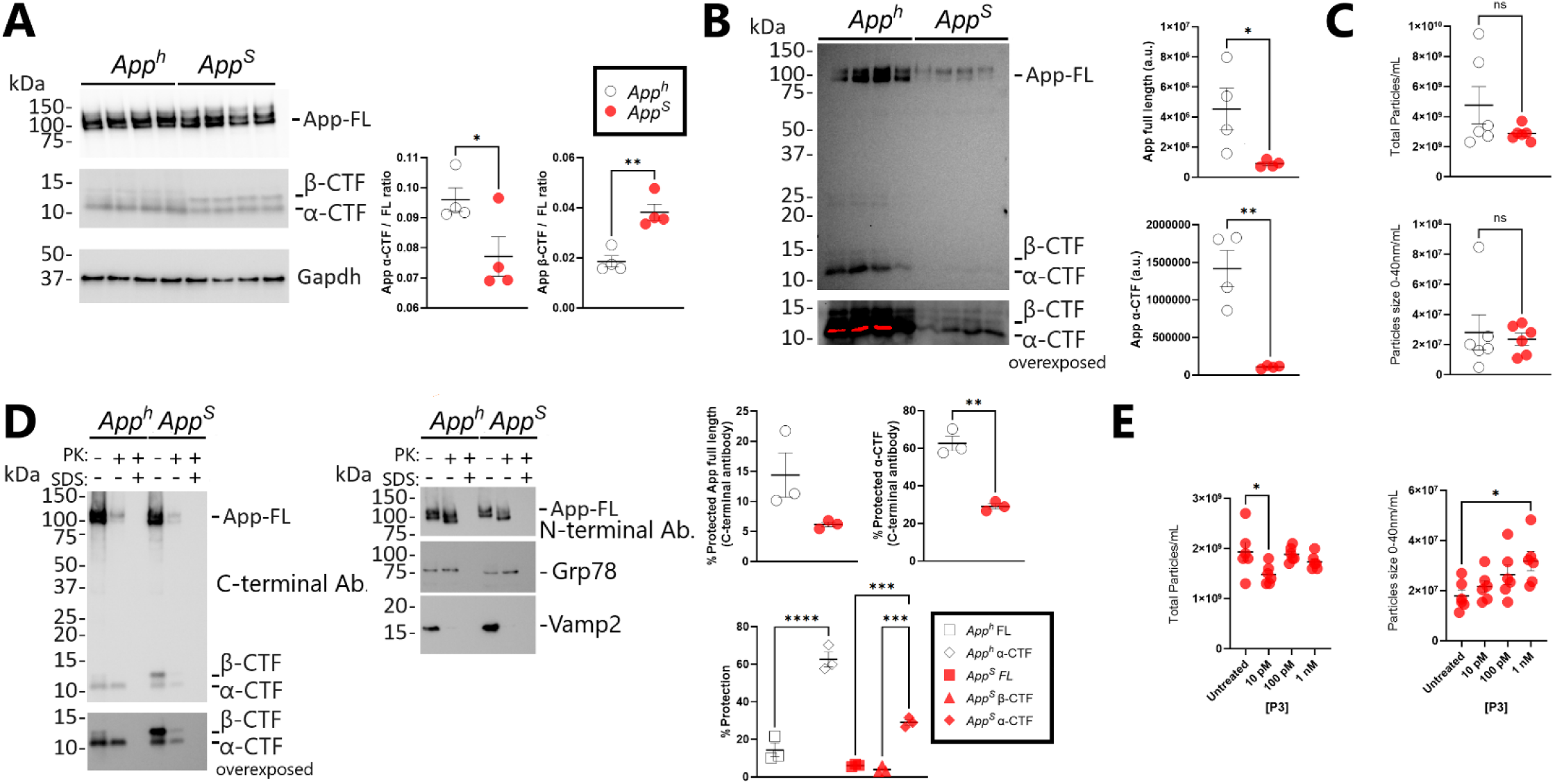
Effect of *App^S^* mutation on App processing and EV levels. **A.** Western analysis of *App^h^* and *App^S^* rat primary neuronal lysate. App full length, App β- and α-CTFs, and Gapdh are indicated. Levels of App α- and β-CTF relative to App full length are represented as mean ± S.E.M. and were analyzed by Student’s *t*-test. \**p* < .05, \*\**p* < .01, n=4. **B.** Western analysis of total EVs isolated from *App^h^* and *App^S^* rat primary neuronal conditioned media, harvested after 24h. App full length, App α- and β-CTF are indicated, with an additional overexposure of App-CTFs shown below. App full length and App α-CTF levels are represented as mean ± S.E.M. and were analyzed by Student’s *t*-test. \**p* < .05, \*\**p* < .01, n=4. **C.** Nanoparticle tracking analysis of conditioned media from *App^h^*and *App^S^* primary neurons. Total particle levels and 0-40 nm particle levels are represented as mean ± S.E.M. and were analyzed by Student’s *t*-test. n=6. **D.** Proteinase protection assay of p0 *App^h^* and *App^S^* rat brain post mitochondrial membranous fractions. Samples were treated with proteinase K (PK) and/or sodium dodecyl sulfate (SDS). App full length and App-CTFs were detected by western analysis with Y188, against App C-terminal epitopes (left), with an additional overexposure of App-CTFs shown below. App full length N-terminal epitopes were detected with 6E10 (right, top). Grp78 (right, middle) and Vamp2 (right, bottom) are indicated. The percentages of PK-protected App metabolites are represented as mean ± S.E.M. and were analyzed by Student’s *t*-test (for *App^h^* and *App^S^* comparisons) or one-way ANOVA (for App metabolite comparisons) with Tukey’s multiple comparison test when ANOVA showed significant differences. \*\**p* < .01; \*\*\**p* < .001, \*\*\*\**p* < .0001, n=3. **E.** Nanoparticle tracking analysis of conditioned media from 24h p3-treated *App^S^* primary neurons. Total particle levels (left) and 0-40 nm particle levels (right) are represented as mean ± S.E.M. and were analyzed by one-way ANOVA with Tukey’s multiple comparison test when ANOVA showed significant differences. \**p* < .05, n=6.

### App α-cleaved neoepitope determines App-EV sorting

The absence of App β-CTF in EVs seen in Fig. 3B may be the result of a retention of App β-CTF-containing ILVs within the cell or, alternatively, a failure of App β-CTF to traffic to ILVs. To distinguish between these two possibilities, a proteinase protection assay was performed. Transmembrane proteins retain their original membrane topology when membranous fractions are homogenized without detergents. Cytosolic-facing regions can be mapped by proteinase K digestion of exposed epitopes. Lumen-facing epitopes are protected from proteinase K digestion by the intact membrane. The C-terminus of a type-I transmembrane protein, such as App, faces the cytosol and is therefore susceptible to proteinase K cleavage. However, if App traffics to a double membranous structure, e.g. to ILVs within MVBs, the C-terminus is protected by the limiting membrane. To eliminate other sources of double membranous vesicles, a post mitochondrial supernatant was prepared from p0 brain homogenates from *App^h^* and *App^S^* rats, treated with proteinase K, and analyzed by western blot. Intralumenal Grp78 and cytosolic Vamp2 controls displayed the expected digestion pattern (**Fig. 3D**, right middle and bottom). Full length App from both *App^h^* and *App^S^* rats displayed the expected proteinase K digestion pattern of a type-I transmembrane protein, with a vast majority of the C-terminal epitopes susceptible to digestion (**Fig. 3D**, left). The N-terminus of App, which is detected by the 6E10 antibody, was protected from degradation but instead shifted down ∼5 kDa in a manner consistent with the digestion of the exposed C-terminus (**Fig. 3D**, right, top). App β-CTF, seen exclusively in *App^S^*homogenates, was also digested in the same pattern as full length App. Interestingly in *App^h^* homogenates, C-terminal epitopes of App α-CTF were mostly protected, indicating that App α-CTF preferentially trafficked to ILVs (**Fig. 3D**, left, bottom).

App α- and β-CTF differ by an additional 16 amino acids present at the N-terminus of App β-CTF, and these N-terminal neoepitopes may underlie the different trafficking pattern of these App-CTFs. The N-terminus of App α-CTF is flexible and loops back into the membrane to terminate at the surface, whereas the additional N-terminal 16 amino acids in App β-CTF re-emerge in the intralumenal/extracellular space^46^. Interestingly, a significantly higher percentage of App α-CTF is protected in *App^h^* samples as compared to *App^S^* samples (**Fig. 3D**), suggesting that the N-terminal neoepitope of App α-CTF alone does not determine ILV/EV localization. P3, the soluble product of App α-CTF digestion by γ-secretase, contains the same α-cleaved neoepitope present in App α-CTF. As p3 is the product of the α- and γ-secretase pathway, the Swedish *App* mutation results in lower p3 production. To determine the contribution of soluble p3 to the decrease in EVs seen in *App^S^* samples, recombinant p3-40 (corresponding to Aβ17-40) was added to *App^S^* primary neuronal cultures, and total EVs were analyzed by nanoparticle tracking. Three doses of p3 were used which span the p3 concentrations found in human CSF^47^. Increasing doses of p3 showed no increase in total EV levels, with a slight but statistically significant decrease found at the 10 pM dose (**Fig. 3E**). When EVs were analyzed by size, a dose-dependent increase was seen in small EV (<40 nm), corresponding to the size of App-EVs (**Fig. 1F**), reaching statistical significance at the 1 nM dose. The neoepitope formed by α-cleavage of App, present in both the membrane bound App α-CTF and soluble p3, may therefore modulate small EV biogenesis.

## METHODS

### Animals

All animal breedings, maintenance, care, and experimental use was performed in accordance with the NIH Guide for the Care and Use of Laboratory Animals. Rutgers Institutional Animal Care and Use Committee has approved the experimental use of animals generated in this study (Protocol #PROTO202200104). Colony genotyping was performed by Transnetyx (TN, USA).

### Primary Neuronal Culture

Plates and flasks were coated overnight with poly-L-lysine (Sigma P4707) and washed 3X with deionized water prior to use. Total cortex was dissected, and meninges were removed from p0-1 rat pup brains. Dissected cortical tissues were digested with trypsin (Gibco 25200056), triturated, filtered with a 0.70 μm cell strainer, and plated onto coverslips or flasks. 12-well plates without coverslips were seeded at 5×10^5^ cells/well for western analysis and nanoparticle tracking analysis. T-75 flasks were seeded at 7.5×10^6^ cells/flask and T-175 flasks were seeded at 1.5×10^7^ cells/flask. Neurons were maintained in Neurobasal (Gibco 21103049) supplemented with 10% B-27 (Gibco 17504044), 1% Pen-Strep (Gibco 15140163), and 2mM glutamine (Gibco 25030081). Cultures were incubated at 37°C and 5% CO_2_ and given half-feeds twice a week. Neurons were treated with 2.5 μM NMS-873 (Sigma SML1128), 25 μM Smer28 (Sigma S8197), 25 μM VCP Activator 1 (MedChemExpress HY-157508), and 50 μM chloroquine (Cell Signaling Technology 14774). Neurons were treated with 0-1 nM p3/Aβ17-40 peptide (Anaspec AS-22813).

### Total EV Isolation

14 DIV primary neuronal conditioned media was collected from 1 T-75 flask per biological replicate, and debris was removed by 0.22 μm PVDF syringe filtration. Total EVs were pelleted by ultracentrifugation of the filtrate at 150,000 × g for 1h at 4°C. Total pellet was lysed in 1X loading buffer (LDS Thermo 84788 supplemented with 10% β-mercaptoethanol) prior to western analysis.

### Immunocapture of App-EVs

21 DIV primary neuronal conditioned media was collected from 4 T-175 flasks per biological replicate, and debris was removed by 0.22 μm PVDF syringe filtration. The filtrate was then concentrated to 80X with a 100 kDa MWCO Vivaspin 20 filter (Sigma Z614661) at 3000 × g, 4°C. Anti-App 4G8 (BioLegend 800703), which targets amino acids 17-24 of Aβ, or Anti-Mouse IgG1 kappa Isotype Control (Thermo 14-4714-82) was bound to Protein A/G agarose beads (Thermo 20421) at 4°C for 1h with end-over-end rotation, at a concentration of 20 μg antibody per 100 μL beads. Unbound antibody was washed off 3X with IP buffer (1 mM EDTA, 50 mM Tris, 150 mM NaCl, pH 8) at 500 × g for 1m. Primary neuronal conditioned media concentrate was incubated with 4G8 or anti-IgG beads at 4°C overnight with end-over-end rotation. Beads were washed 7X with IP buffer and eluted with 300 ng/mL Aβ17-24 peptide (Anaspec AS-61978) for 1h at room temperature with gentle agitation. Eluted EVs were used for downstream mass spectrometry, western, or transmission electron microscopic analysis.

### Proteinase K Protection Assay

Brains from p0 pups were homogenized with a glass-glass homogenizer in SEMK buffer (220 mM sucrose, 10 mM MOPS, 1 mM EDTA, 20 mM KCl, pH 7.2), supplemented with 1% protease/phosphatase inhibitor cocktail (Sigma PPC1010), on ice. The homogenate was centrifuged twice at 15,000 × g for 10m to produce a post-mitochondrial supernatant. The supernatant was ultracentrifuged at 150,000 × g for 1h to produce a pellet containing membranous organelles. The pellet was resuspended in SEMK buffer and total protein content was determined by Bradford analysis. In a 50 μL reaction, 50 μg of the membranous organellar fraction in SEMK buffer was digested with 0.5 μL of 2 μg/mL proteinase K (PK) (Sigma P6556) at 37°C for 10m. A negative control without proteinase K and a positive control with proteinase K and 0.1% SDS were performed simultaneously. Digestion was halted with 100 mM phenylmethylsulfonyl fluoride (Roche 10837091001) and by boiling reaction mixture for 10m at 100°C. Total reaction mixture was analyzed by western analysis.

### Mass Spectrometry

Mass spectrometry experiments were performed by MSBioworks (MI, USA) as follows: App-EV eluate was processed by SDS-PAGE using a 10% Bis-Tris NuPAGE gel (Invitrogen) with the MES buffer system. The mobility region was excised into 10 equal sized segments and in-gel digestion was performed on each using a robot (DigestPro, CEM) with the following protocol: washed with 25 mM ammonium bicarbonate followed by acetonitrile, reduced with 10 mM dithiothreitol at 60°C followed by alkylation with 50 mM iodoacetamide at RT, digested with sequencing grade trypsin (Promega) at 37°C for 4h, and quenched with formic acid. The supernatants were combined and lyophilized. Samples were dissolved in 0.1% TFA for analysis.

Half of each digested sample was analyzed by nano LC-MS/MS with a Waters M-Class LC system interfaced to a ThermoFisher Exploris 480 mass spectrometer. Peptides were loaded on a trapping column and eluted over a 75 μm analytical column at 350 nL/min; both columns were packed with XSelect CSH C18 resin (Waters); the trapping column contained a 3.5 μm particle, the analytical column contained a 2.4 μm particle.

The column was heated to 55°C using a column heater (Sonation). The mass spectrometer was operated in data-dependent mode, with the Orbitrap operating at 60,000 FWHM and 15,000 FWHM for MS and MS/MS respectively. The instrument was run with a 3s cycle for MS and MS/MS. Advanced Precursor Determination^48^ was enabled. 5h of instrument time was used for the analysis of each sample.

Data were searched using a local copy of Mascot (Matrix Science) with the following parameters: Enzyme: Trypsin/P; Database: UniProt Rat (concatenated forward and reverse plus common contaminants); Fixed modification: Carbamidomethyl (C); Variable modifications: Oxidation (M), Acetyl (N-term), Pyro-Glu (N-term Q), Deamidation (N,Q); Mass values: Monoisotopic; Peptide Mass Tolerance: 10 ppm; Fragment Mass Tolerance: 0.02 Da; Max Missed Cleavages: 2. Mascot DAT files were parsed using Scaffold (Proteome Software) for validation, filtering and to create a non-redundant list per sample. Data were filtered at 1% protein and peptide FDR and requiring at least two unique peptides per protein.

### Cell Surface Labeling

Total primary neuronal surface membrane proteins were labeled with Sulfo-NHS-SS-biotin and isolated by immunoprecipitation adapted from published protocols^49^. Briefly, primary neurons grown in 12-well plates were biotinylated with 0.3 mL of 0.5 mg/mL Sulfo-NHS-SS-biotin solution (Thermo 21331) for 30m on ice. Unreacted linker was quenched with 50 mM glycine in PBS 3X for 5m on ice. Neurons were lysed in 120 μL IP buffer supplemented with 1% Triton-X for 10m on ice. Lysate was centrifuged at 17,000 x g for 10m at 4°C and the supernatant was collected. An aliquot of total lysate was stored separately for western analysis. Total surface protein was isolated by immunoprecipitation with Neutravidin beads (Thermo 29200). 50 μL of 50% bead slurry was incubated with 100 μL lysate for 2h at 4°C with end-over-end rotation. Beads were washed 7X with IP buffer and total surface protein was eluted by boiling for 1m in 50 μL 1X loading buffer.

### Aβ ELISA

Primary neuronal conditioned media from 14 DIV neurons was collected and dead cells were removed by 0.22 μm PVDF syringe filtration. Conditioned media levels of Aβ40 and Aβ42 were determined by Meso Scale Discovery (MSD) multi-array electrochemiluminescence assay kit (K15199G-1). MSD kit was used according to manufacturer’s recommendations and read on a MESO QuickPlex SQ 120 plate reader.

### Western Analysis

For analysis of conditioned media, total conditioned media was passed through a 0.22 μm PVDF syringe filter. 1X loading buffer was added to the filtrate, which was then boiled for 1m and loaded. For analysis of cell lysate, primary neurons were lysed in RIPA buffer (10 mM Tris-HCl, pH 8.0, 1 mM EDTA,1% Triton X-100, 0.1% SDS, 140 mM NaCl). 15 μg of protein was brought to 15 μl with PBS and 1X loading buffer and loaded on a 4%–12% BisTris polyacrylamide gel (Bio-Rad 3450125). Proteins were transferred onto nitrocellulose at 25 V for 7m using the Trans-blot Turbo system (Bio-Rad) and visualized by red Ponceau staining. Membranes were blocked for 1h in 5% milk (Bio-Rad 1706404) and washed extensively in PBS/Tween 20 (0.05%). Primary antibody was applied overnight at 4°C at 1:1000 dilution in 5% BSA (Fisher BP9703100). The following primary antibodies were used: 6E10 used for sAppα (App Aβ3–8 epitope, Biolegend 803001), Y188 used for App (App-C-terminus epitope, Abcam AB32136), Gapdh (Cell Signaling Technology 2118), Alix (Cell Signaling Technology 92880), Vcp (Cell Signaling Technology 2649), Lc3b (Cell Signaling Technology 83506), flotillin-1 (Cell Signaling Technology 18634), CD9 (Cell Signaling Technology 98327), Bip/Grp78 (Cell Signaling Technology 3183), and Vamp2 (Synaptic Systems 104202). Primary antibodies were washed off extensively and 1:1000 dilutions of secondary antibodies, either anti-mouse (Southern Biotech 1030-05) or anti-rabbit (Southern Biotech 4030-05) in 5% milk PBS/Tween 20, were applied for 1h at room temperature with shaking. Blots were developed with Clarity and Clarity Max ECL Western Blotting Substrates (Bio-Rad 1705060 and 1705062) and visualized on a ChemiDoc MP Imaging System (Bio-Rad). Signal intensity was quantified with Image Lab software (Bio-Rad).

### Immunoprecipitation

Total p0 brain lysate was diluted in IP buffer supplemented with 1% Triton-X100, solubilized for 1h at 4°C with end-over-end rotation. Samples were spun at 17,000 × g for 10m. Solubilized lysate was used as input for immunoprecipitation with anti-App 4G8, anti-Vcp (Invitrogen MA3-004), or control anti-Mouse IgG1 kappa Isotype Control and protein A/G beads overnight at 4 °C with end-over-end rotation. Beads were washed 7X with IP buffer, and bound protein was eluted by 1m boiling in 1X loading buffer. Input (diluted 1:20 in 1X loading buffer) and eluates were analyzed by western blot analysis.

### Statistical Analysis

Statistical significance was evaluated using ordinary one-way ANOVA followed by post hoc Tukey’s multiple comparisons test when applicable (*i.e.* when the ordinary one-way ANOVA showed statistical significance) or by Student’s *t*-test. Statistical analysis was performed with GraphPad Prism v10 for Windows. Significant differences were accepted at *p* < 0.05, with error bars representing SEM.

### Nanoparticle Tracking Analysis

Conditioned primary neuronal culture media was analyzed by Alpha Nano Tech (Morrisville, NC). Briefly, samples were diluted with fresh 0.2 μm filtered (Sarstedt 831826001) deionized water to achieve a concentration of 100-300 particles per screen. The diluted samples were briefly vortexed and loaded into 1 mL syringes for loading into the machine. Zetaview Quatt NTA instrument (Particle Metrix, Meerbusch, Germany) was used for analyzing after alignment with 100 nm polystyrene beads. The following instrument settings were used: Mode at Scatter (488 nm), Sensitivity at 83, Shutter at 100, Cycles/positions at 1/11, Frame rate at 30, Maximum Size at 1000, Minimum Size at 20, Track Length at 15, Minimum Brightness at 20. Data in figures are represented after dilution factor adjustments.

### TEM

Isolated App-EVs were analyzed by Alpha Nano Tech. Briefly, copper carbon Formvar grids were cleaned with glow discharge and floated on a sample drop for 10 minutes for sample adsorption. The grids were then washed twice by floating on a drop of deionized water and stained with 2% uranyl acetate for imaging using JEM-1230 (Jeol).

## DISCUSSION

This study (1) identifies Vcp as molecular cargo in App-EVs, (2) explores the consequence of amyloidogenic vs nonamyloidogenic processing of App in App-EV biogenesis, and (3) uncovers a new role of Vcp in EV secretion. There is a significant genetic connection between Vcp and neurodegeneration. Autosomal dominant mutation of *VCP* causes tau-only frontotemporal dementia^50^, multisystem proteinopathy^51^, characterized by frontotemporal dementia with tauopathy plus extra-CNS proteinopathies in muscle and bone, and amyotrophic lateral sclerosis^52^, characterized by upper motor neuron loss and intracellular proteinopathy. No *VCP* mutation has been found to cause AD, though some observations support a functional link between VCP and the main pathological features of AD, i.e. tau and amyloid. The VCP-tau connection has been established by numerous studies which show that VCP can directly bind aggregated tau, disassemble it, and potentially affect the spread of tau tangle pathology^50,53,54^. Evidence of a relationship between VCP function and amyloid is less established. Patients with inclusion-body myositis, related to the muscle proteinopathy caused by *VCP* mutations, have amyloid positive rimmed vacuoles within muscle cells^55,56^. Additionally, in AD patients, VCP is increased in brain-derived EVs, as compared to nondemented controls^57^. Our study is the first to provide a direct cell biological link between Vcp and amyloid in the form of Vcp as molecular cargo in App-EVs. The ability of Vcp to localize to App-EVs suggests a new cellular function of App related to the clearance of protein aggregates.

The precise cellular function of APP is unknown. Most studies of AD-causing *APP* mutations focus on the biochemical changes to Aβ amount, Aβ length, and Aβ self-association. These processes affect the extent to which Aβ aggregates, and this metric features heavily in how AD is defined. However, Aβ is just one metabolite of APP, and the mutations which govern Aβ production also affect the numerous non-Aβ metabolites as well as the function of the full length protein. Altered APP function may have a pleiotropic effect: one manifestation of which is the production of aggregation-prone Aβ species, and another contemporaneous manifestation is a change in APP-EV production. The advantage of expanding our understanding of APP function to include APP-EVs is that it allows for a new connection between amyloid and tau. Vcp, in its capacity to disaggregate tau and localize to App-EVs, may underpin this link between the two canonical AD pathologies.

All *APP* mutations that cause or prevent AD occur within the juxtamembranous or transmembrane regions of APP^3^. These regions are also the subject of extensive proteolysis which results in different species of APP-CTFs, with APP α- and β-CTFs most abundant. Unexpectedly, we have found that this region governs the localization of App-CTFs to ILVs, with App α-CTF localizing predominantly to ILVs while the longer App β-CTF and full length App are only present in ILVs in minor amounts (**Fig. 3D**). The biophysical cause of this change in ILV localization is unclear but may be related to changes in membrane curvature that are required for the invagination of the limiting endosomal membrane to form ILVs. The juxtamembranous region of App α-CTF re-inserts into the endosomal membrane^46^. This close apposition may modulate membrane curvature and be lost when longer or mutated forms of App-CTFs are present. This ability may not be limited to membrane-bound forms of App. We also considered the effect of soluble forms of App which contain this neoepitope formed by α-cleavage, such as p3, and found that exogenous p3 increases small EV biogenesis. These observations support the further study of the numerous *APP* mutations which modulate the composition or abundance of APP-CTFs or p3.

The study of App-EVs has uncovered a new function of Vcp in its ability to cause global secretion of EVs. This new function is relevant to AD and other types of neurodegeneration, as EVs have been proposed as a mechanism for the cell-to-cell spread of toxic protein aggregates^58^, including tau. We speculate that the disruption of the autophagy-promoting ability of Vcp, accomplished pharmacologically in our study, mimics the autophagy failure seen in AD patients and animal models. Secretion of EVs may be an alternative route of clearance when normal degradative pathways are impaired. Therefore, in addition to the known effects of Vcp function on the seeding-potential of tau aggregates *within* a cell^53^, Vcp function may be relevant for the *cell-to-cell* spread of tau as well.

## ACKNOWLEDGEMENTS

Nanoparticle tracking analysis and electron microscopy were performed by Alpha Nano Tech (Morrisville, NC). Colony genotyping was performed by Transnetyx (TN, USA). Mass spectrometry experiments were performed by MSBioworks (MI, USA).

This work was supported by NIA, National Institutes of Health Grant R00AG065441 (to M.D.T.). The authors declare that they have no conflicts of interest with the contents of this article. The content is solely the responsibility of the authors and does not necessarily represent the official views of the National Institutes of Health.

## COMPETING INTERESTS

The authors declare no competing interests.

---

**Figure S1.**
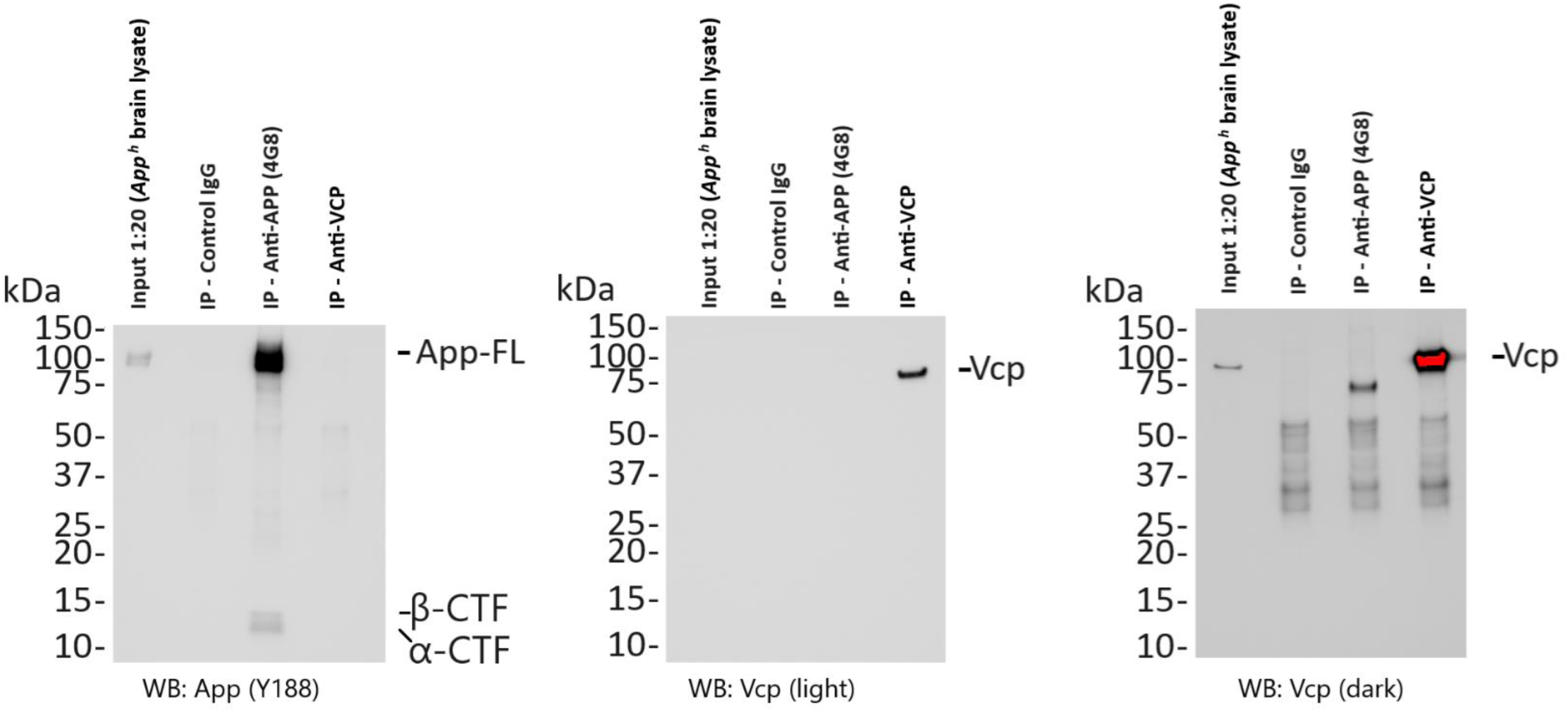
Co-immunoprecipitation of App and Vcp in *App^h^* total brain lysate. App and Vcp were immunoprecipitated from total brain lysate, with 4G8 and MA3-004, respectively. Eluate was analyzed by western blot against App (Y188) and Vcp (CST 2649).

**Figure S2.**
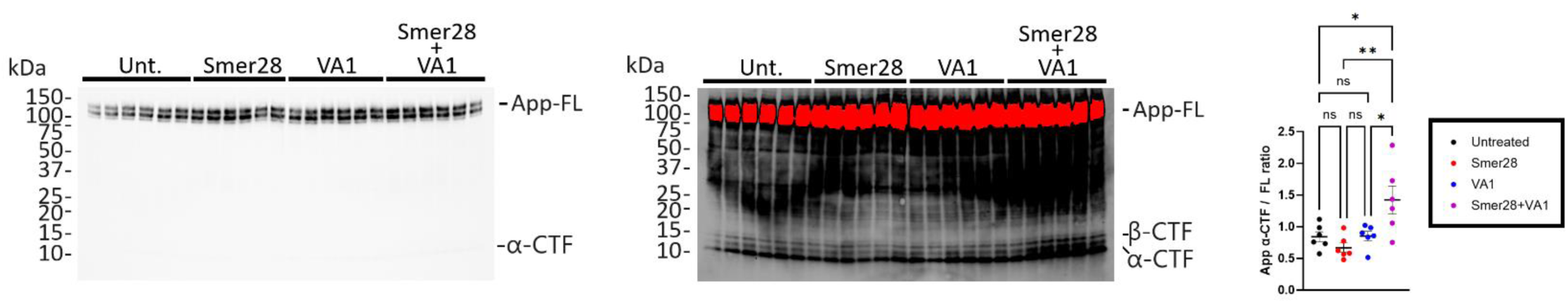
Effect of Vcp activation on App α-CTF levels. Western analysis of lysate from *App^h^*rat primary neurons treated with Smer28 and/or VA1 for 24h. App full length (left) and App-CTFs (middle) are indicated. App α-CTF levels relative to App full length levels (right) are represented as mean ± S.E.M. and were analyzed by one-way ANOVA with Tukey’s multiple comparison test when ANOVA showed significant differences. \**p* < .05, \*\**p* < .01, n=6.

## Notes

### Competing Interest Statement

The authors have declared no competing interest.

